# Boosting with ALVAC-HIV and AIDSVAX B/E enhances Env constant region 1 and 2 antibody-dependent cellular cytotoxicity

**DOI:** 10.1101/632844

**Authors:** David Easterhoff, Justin Pollara, Kan Luo, William D. Tolbert, Brianna Young, Dieter Mielke, Shalini Jha, Robert J. O’Connell, Sandhya Vasan, Jerome Kim, Nelson L. Michael, Jean-Louis Excler, Merlin L. Robb, Supachai Rerks-Ngarm, Jaranit Kaewkungwal, Punnee Pitisuttithum, Sorachai Nitayaphan, Faruk Sinangil, James Tartaglia, Sanjay Phogat, Thomas B. Kepler, S. Munir Alam, Kevin Wiehe, Kevin O. Saunders, David C. Montefiori, Georgia D. Tomaras, M. Anthony Moody, Marzena Pazgier, Barton F. Haynes, Guido Ferrari

## Abstract

Induction of protective antibodies is a critical goal of HIV-1 vaccine development. One strategy is to induce non-neutralizing antibodies that kill virus-infected cells as these antibody specificities have been implicated in slowing HIV-1 disease progression and in protection. HIV-1 Env constant region 1 and 2 (C1C2) antibodies frequently contain potent antibody dependent cellular cytotoxicity (ADCC) making them a vaccine target. Here we explore the effect of delayed and repetitive boosting of RV144 vaccinee recipients with ALVAC/AIDSVAX B/E on the C1C2-specific antibody repertoire. It was found that boosting increased clonal lineage specific ADCC breadth and potency. A ligand crystal structure of a vaccine-induced broad and potent ADCC-mediating C1C2-specific antibody showed that it bound a highly conserved Env gp120 epitope. Thus, rationally designed boosting strategies to affinity mature these type of IgG C1C2-specific antibody responses may be one method by which to make an improved HIV vaccine with higher efficacy than seen in the RV144 trial.

**Significance:** Over one million people become infected with HIV-1 each year making the development of an efficacious HIV-1 vaccine an important unmet medical need. The RV144 human HIV-1 vaccine-regimen is the only HIV-1 clinical trial to date to demonstrate vaccine-efficacy. An area of focus has been on identifying ways by which to improve upon RV144 vaccine-efficacy. The RV305 HIV-1 vaccine-regimen was a follow-up boost of RV144 vaccine-recipients that occurred 6-8 years after the conclusion of RV144. Our studies focused on the effect of delayed boosting in humans on the vaccine-induced antibody repertoire. It was found that boosting with a HIV-1 Env vaccine increased antibody-mediated effector function potency and breadth.

## INTRODUCTION

CD4-inducible (CD4i) epitopes within HIV-1 envelope (Env) constant regions 1 and 2 (C1C2) are targets for antibodies that mediate antibody dependent cellular cytotoxicity (ADCC) [1]. C1C2-specific antibody epitopes have been termed Cluster A [1] and defined by two Env-targeted monoclonal antibodies (mAbs), A32 [2] and C11 [1]. Structural analyses of antigen complexes formed by A32, A32-like [3–5] and C11-like antibodies [6] indicate that these antibodies bind distinct Env epitopes. The A32 epitope involves a discontinuous sequence within layers 1 and 2 of the inner domain [4, 5] while the C11 epitope maps to the inner domain eight-stranded β sandwich [6]. Importantly, both antibodies are non-neutralizing for tier 2 HIV strains, but are capable of broad and potent ADCC [1, 2].

The secondary analysis of HIV-1 infection risk in RV114 (NCT00223080) indicated that ADCC in the presence of low anti-Env IgA responses correlated with decreased HIV-1 acquisition [7]. While antibodies representative of the Env variable region 2 (V2) response inversely correlated with HIV-1 acquisition [7], we previously demonstrated that synergy between A32-blockable C1C2-specific antibodies and V2-specific antibodies increased the potency of V2 antibodies induced in the RV144 trial [8].

Here we have studied the effects of late boosting of RV144 vaccinees on affinity maturation and potency of C1C2-specific ADCC antibodies in the RV305 HIV-1 vaccine trial (NCT01435135). We found that ALVAC/AIDSVAX B/E immunizations induced C1C2-specific antibodies and that late booster immunizations increased C1C2-specific antibody variable heavy and variable light (VH + VL) chain gene mutation frequencies and increased their ADCC breadth and potency.

## RESULTS

### AIDSVAX B/E N-terminal deletion alters C1C2-specific antibody responses

AIDSVAX B/E protein used in the RV144 and RV305 HIV-1 vaccine trial had an eleven amino acid N-terminal deletion [9] that removed a majority of the C11-like antibody epitope [6], whereas CRF_01 AE gp140 Env 92TH023 in ALVAC (vCP1521) did have the gp120 N-terminal 11 amino acids [10]. To determine if C11 could bind to gp120 proteins with an 11 amino acid N-terminal deletion, we assayed A32 and C11 antibodies for binding to full length AE.A244gp120 or to AE.A244gp120Δ11 (N-terminal 11 aa deleted). Antibody A32 bound to full length AE.A244gp120 and A32 binding was enhanced on AE.A244gp120Δ11 (**Fig. 1A**) [9]. In contrast, antibody C11 only bound to the full length AE.A244gp120 (**Fig 1A**). From these data we concluded that C11-like antibody responses were unlikely to be boosted by AIDSVAX B/E.

**Figure 1.**
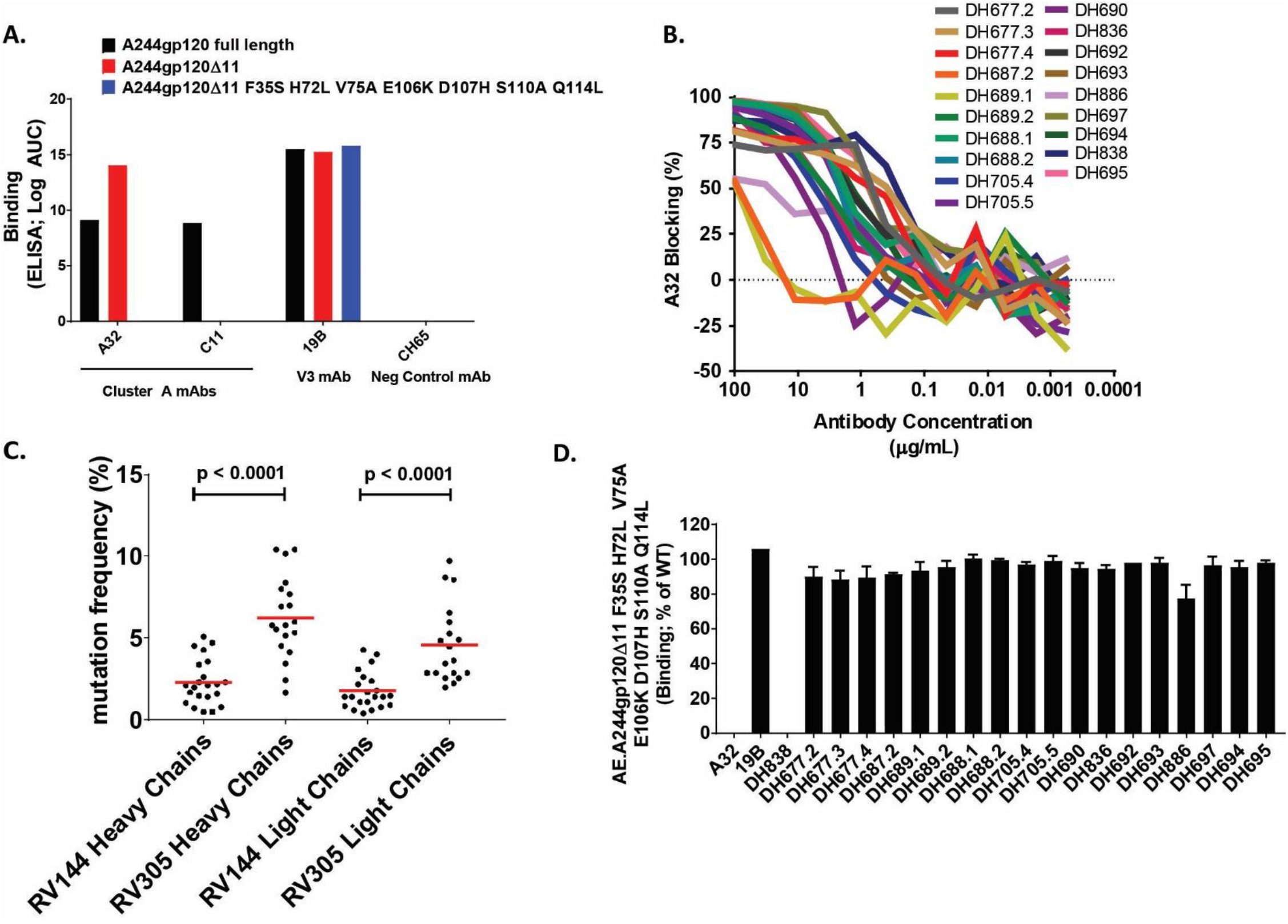
Identification of RV305 C1C2-specific antibodies. (A) The C1C2-specific antibodies A32 or C11 were assayed by direct binding ELISA for reactivity with full length AE.A244gp120 or AE.A244g120Δ11. An A32-specific mutant protein was designed (AE.A244g120Δ11 F35S H72L V75A E106K D107H S110A Q114L) to identify A32-like antibody responses. 19B antibody was used as a positive control and CH65 as a negative control. (B) RV305 non-neutralizing antibodies were assayed for A32-blocking by ELISA. (C) RV305 non-neutralizing A32-blockable antibody heavy and light chain gene sequence mutation frequencies were analyzed by Cloanalyst (Kepler et al., 2014) and compared to previously published RV144 heavy and light chain gene sequence mutation frequencies (% nucleotide) (Bonsignori et al., 2012). Statistical significance was determined using a Wilcoxon rank sum test. Red bar represents that mean (D) RV305 non-neutralizing A32-blockable antibody were assayed by direct binding ELISA to AE.A244g120Δ11 and AE.A244g120Δ11 F35S H72L V75A E106K D107H S110A Q114L. Data are expressed as % binding the mutant protein relative to WT. Shown are the mean with standard deviation of two independent experiments.

A total of 19 RV305-derived NNAbs isolated from four individuals (**Table S1 & S2**) were identified that blocked the C1C2 mAb A32 binding to AE.A244gp120Δ11 (**Fig 1B**). Compared to previously published RV144 C1C2-specific antibodies [11] the RV305 C1C2-specific antibodies had significantly more V_H_ and V_L_ chain gene mutations (Wilcoxon rank sum test P < 0.0001) (**Fig 1C**) suggesting that RV305 boosting induced additional somatic mutations in C1C2-specific antibodies.

To determine if RV305 boosted A32 blockable antibodies contained a binding epitope similar to A32, we used the A32 ligand crystal structure [5] to identify critical A32 antibody contact residues, and then designed an AE.A244gp120Δ11 mutant protein (AE.A244gp120Δ11 F35S, H72L, V75A, E106K, D107H, S110A, Q114L) to eliminate A32-like antibody binding (**Fig 1A**). In ELISA, the RV305 antibody, DH838, was the only antibody with binding eliminated by mutating the A32 epitope (**Fig 1D**). Likewise, DH838 was the only antibody that used a VH3 family gene while all other ALVAC/AIDSVAX B/E – induced C1C2-specific antibodies used VH1 genes (**Table S1**). Thus, as in RV144, ALVAC/AIDSVAX B/E boosting preferentially expanded VH1 gene C1C2-specific antibodies [11] and these antibodies bound epitopes distinct from A32 but in close enough proximity to be sterically cross-blocked by A32 (**Fig 1B**).

### Boosting increased C1C2-specific ADCC breadth and potency

RV305 C1C2-specific antibodies and a subset of RV144 C1C2-specific antibodies were next assessed for ADCC against a cross-clade panel of HIV-1 infectious molecular clone (IMC) infected CD4+ T cells (HIV-1 AE.CM235, B.WITO, C.TV-1. C.MW965, C.1086C, C.DU151 and C.DU422). Antibodies were ranked using an ADCC score (See methods) that accounted for ADCC breadth and potency. Apart from the RV144-derived A32 blockable antibody CH38, which was naturally an IgA antibody but tested here as a recombinant IgG1 antibody, 16/19 RV305 antibodies ranked higher than the RV144 antibodies (**Table 1**). Next RV305 derived C1C2-specific antibody heavy chain gene mutation frequency was used as a proxy for responsiveness to boosting and compared to the ADCC score. The V_H_ mutation frequency (% nucleotide) inversely correlated with the ADCC score (Spearman Correlation −0.5599; p value = 0.0127) (**Fig S1**). However the V_H_ mutation frequency of those antibodies with the highest ADCC scores were above the average heavy chain gene mutation frequency for RV144 (**Fig S1 Fig 1C**). Thus, while a RV144 boosting regimen was necessary to increase C1C2-specific ADCC breadth and potency, additional boosting with same immunogens may not be beneficial.

**Table 1:** Ranking C1C2-specific antibodies by ADCC breadth and potency. RV305 and RV144 C1C2-specificantibodies were assayed for antibody-dependent cellular cytotoxicity against AE.CM235, B.WITO, C.TV-1, C.MW965, C.1086C, C.DU151 and C.DU422 infectious molecular clone infected cells. Antibodies were ranked using an ADCC Score that accounts for breadth and potency (see methods). Number of strains recognized was determined by ADCC endpoint concentration > 40μg/mL.

| **Rank** | **Antibody** | **Study** | **Score** | **Average AUC** | **Number of strains recognized** |
| --- | --- | --- | --- | --- | --- |
| **1** | A32 |  | 6.62 | 198.70 | 7 |
| **2** | CH38 | RV144 | 6.28 | 186.56 | 6 |
| **3** | DH677.3 | RV305 | 4.56 | 146.90 | 7 |
| **4** | DH697 | RV305 | 2.70 | 111.98 | 7 |
| **5** | DH677.2 | RV305 | 1.72 | 111.76 | 5 |
| **6** | DH838 | RV305 | 1.62 | 107.24 | 4 |
| **7** | DH677.4 | RV305 | 1.30 | 94.00 | 6 |
| **8** | DH695 | RV305 | 0.86 | 82.94 | 7 |
| **9** | DH688.1 | RV305 | 0.66 | 77.94 | 6 |
| **10** | DH694 | RV305 | 0.58 | 80.18 | 5 |
| **11** | DH705.5 | RV305 | 0.52 | 75.84 | 5 |
| **12** | DH705.4 | RV305 | 0.46 | 75.46 | 6 |
| **13** | DH690 | RV305 | 0.36 | 67.44 | 7 |
| **14** | DH692 | RV305 | 0.08 | 65.14 | 6 |
| **15** | DH693 | RV305 | 0.06 | 64.54 | 7 |
| **16** | DH886 | RV305 | -0.08 | 67.96 | 5 |
| **17** | DH688.2 | RV305 | -0.30 | 60.42 | 5 |
| **18** | DH836 | RV305 | -0.32 | 62.52 | 5 |
| **19** | CH57 | RV144 | -0.48 | 53.22 | 5 |
| **20** | CH90 | RV144 | -1.06 | 38.98 | 5 |
| **21** | CH54 | RV144 | -1.20 | 34.58 | 5 |
| **22** | DH689.2 | RV305 | -1.42 | 24.54 | 6 |
| **23** | DH689.1 | RV305 | -1.68 | 26.48 | 6 |
| **24** | DH677.1 | RV144 | -2.20 | 16.38 | 2 |
| **25** | DH687.2 | RV305 | -2.70 | 0.16 | 2 |

### Boosting of RV144 vaccinees in the RV305 trial increased ADCC breadth and potency of the RV144 derived C1C2-specific, DH677 clonal lineage

Next the C1C2-specific DH677 memory B cell clonal lineage was used to study affinity maturation and ontogeny of ALVAC/AIDSVAX B/E-induced ADCC responses. B cell clonal lineage member DH677.1 arose after the original RV144 trial (ALVAC + AIDSVAX B/E) and the DH677.2, DH677.3 and DH677.4 clonal lineage members were isolated after delayed and repetitive boosting with AIDSVAX B/E alone (RV305 Group II). Thus, this B cell clonal lineage belongs to a long-lived memory B cell pool started by the RV144 vaccine-regimen and boosted many years later with the RV305 vaccine-regimen (**Fig 2**). The DH677 clonal lineage was assayed by surface plasmon resonance for binding to the AIDSVAX B/E proteins – AE.A244gp120Δ11 and B.MNgp120Δ11 – as well as full length AE.A244gp120. DH677 unmutated common ancestor (UCA) did not bind to B. MNgp120Δ11, had minimal binding to the full length AE.A244gp120 and this binding was enhanced with AE.A244gp120Δ11 (**Fig 2 and Fig S2**). The RV305 boosts more than doubled the V_H_ chain gene mutation frequency from 1.04% (DH677.1; RV144) up to 4.51% (DH677.4; RV305) which resulted in 100-fold increase in apparent affinity for the AIDSVAX B/E proteins (DH677.1 AE.A244gp120Δ11 K_D_= 45.2 & B.MNgp120Δ11 K_D_ =219 to DH677.4 AE.A244gp120Δ11 K_D_= 0.49 & B.MNgp120Δ11 K_D_ =2.86) and also improved binding to full length AE.A244gp120 (**Fig 2 and Fig S2**).

**Figure 2.**
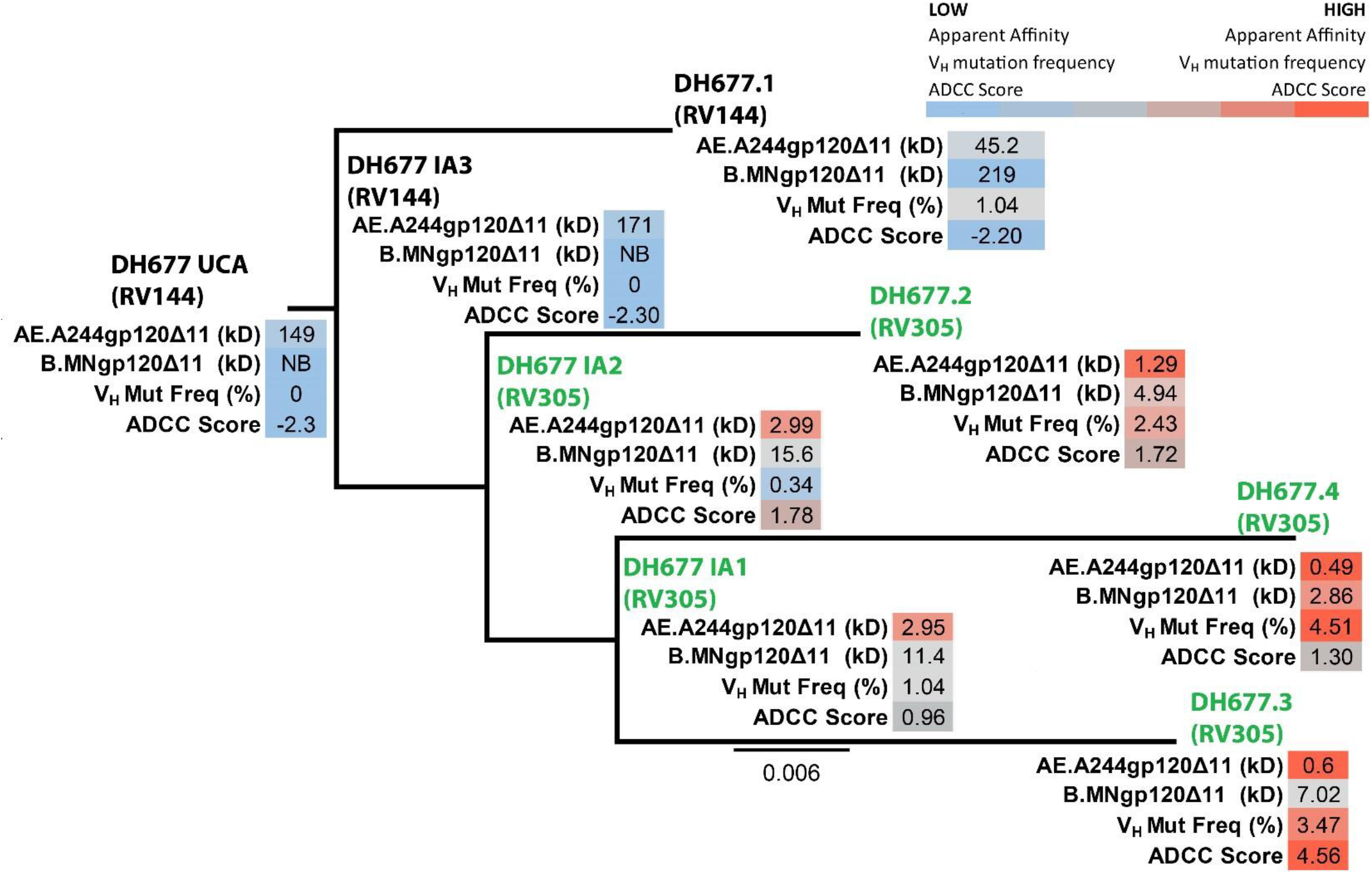
RV305 boosting increased the apparent affinity and antibody dependent cellular cytotoxicity breadth and potency of the C1C2-specific RV144 derived DH677 memory B cell clonal lineage. DH677.1 antibody was isolated by antigen-specific single-cell sorting PBMC from the RV144 vaccine trial. DH677.2, DH677.3 and DH677.4 were isolated by antigen-specific single-cell sorting PBMC collected after the second boost AIDSVAX B/E (RV305 Group II) given in RV305 (~7yrs later). The intermediate ancestors and unmutated common ancestor was inferred using Clonalayst [31]. Recombinantly expressed antibodies were assayed by biolayer interferometry for binding to the AIDSVAX B/E proteins – AE.A244g120Δ11 + B.MNg120Δ11- and for antibody dependent cellular cytotoxicity (ADCC) against AE.C235, B.WITO, C.TV-1, C.MW965, C.1086C, C.DU151 and C.DU422 Renilla luciferase reporter gene infectious molecular clone infected cells. An ADCC score (see methods) was used to account for breadth and potency.

The ontogeny of vaccine-induced ADCC was studied by assaying the DH677 clonal lineage against a cross-clade panel of IMC infected cells (AE.CM235, B.WITO, C. TV-1, C.MW965, C.1086C, C.DU151 and C.DU422). The RV144 prime-boost immunization regimen minimally increased ADCC breadth and potency (DH677 UCA ADCC Score = −2.32; DH677.1 ADCC Score = −2.20 (see methods)). Conversely, RV305 boosting substantially increased ADCC breadth and potency (DH677.3 ADCC Score = 4.56) (**Fig 2 and Fig S3**). These data indicated that the RV144 prime-boost regimen was insufficient to fully affinity mature this C1C2-specific B cell clonal lineage. Rather RV305 trial boosting of this particular RV144 vaccinee profoundly enhanced DH677 lineage ADCC breadth and potency.

### Crystal structure of the potent ADCC-mediating antibody DH677.3

We next determined the crystal structures of the antigen binding fragment (Fab) of the highest ranking RV305 ADCC antibody DH677.3 (**Table 1**) – alone and in complex with clade AE gp120_93TH057_ core_e_ plus the CD4-mimetic M48-U1 (**Table S3**). DH677.3 Fab-gp120_93TH057_ core_e_-M48U1 complex (**Fig 3**) showed that, similar to other Cluster A antibodies, DH677.3 approaches gp120 at the face that is buried in the native Env trimer [3–5] and binds the C1C2 region exclusively within the gp120 inner domain. gp120 residues involved in DH677.3 binding map to the base of the 7-stranded β-sandwich (residues 82, 84, 86-87, 222-224, 244-246, and 491-492) and its extensions into the mobile layers 1 (residues 53, 60, 70-80) and 2 (residues 218-221). By docking at the layer 1/2/ β -sandwich junction the antibody buried surface area (BSA) utilizes 248 Å^2^ of the β-sandwich, 542 Å of layer 1 and 135 Å^2^ of layer 2 (Table S4). The majority of contacts providing specificity involve a network of hydrogen bonds and a salt bridge (**Fig 3A, inset**) contributed by the antibody heavy chain and gp120 side chain atoms of layer 1 (α turn connecting the 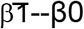 strands, D^78^ and N^80^) and the 7-stranded-β-sandwich (strand β7, Q^246^). The contacts provided by the light chain are less specific and consist of hydrogen bonds to the gp120 main chain atoms and hydrophobic contacts within a hydrophobic cleft formed at the layer 1/2/β-sandwich junction (**Fig 3B and C**). Overall DH677.3 utilizes all six of its complementary determining regions (CDRs), and relies approximately equally on both heavy chain and light chain with a total buried surface area (BSA) of 973 Å^2^: 498 Å^2^ for the light chain and 475 Å^2^ for the heavy chain (**Table S4** and **Fig 3C and S5**). Interestingly, 25 of 29 gp120 contact residues are conserved in >80% of sequences in the HIV Sequence Database Compendium (https://www.hiv.lanl.gov/content/sequence/HIV/COMPENDIUM/compendium.html) with 15 of 29 being effectively invariant (>99% conserved) (**Fig 3B**).

**Figure 3.**
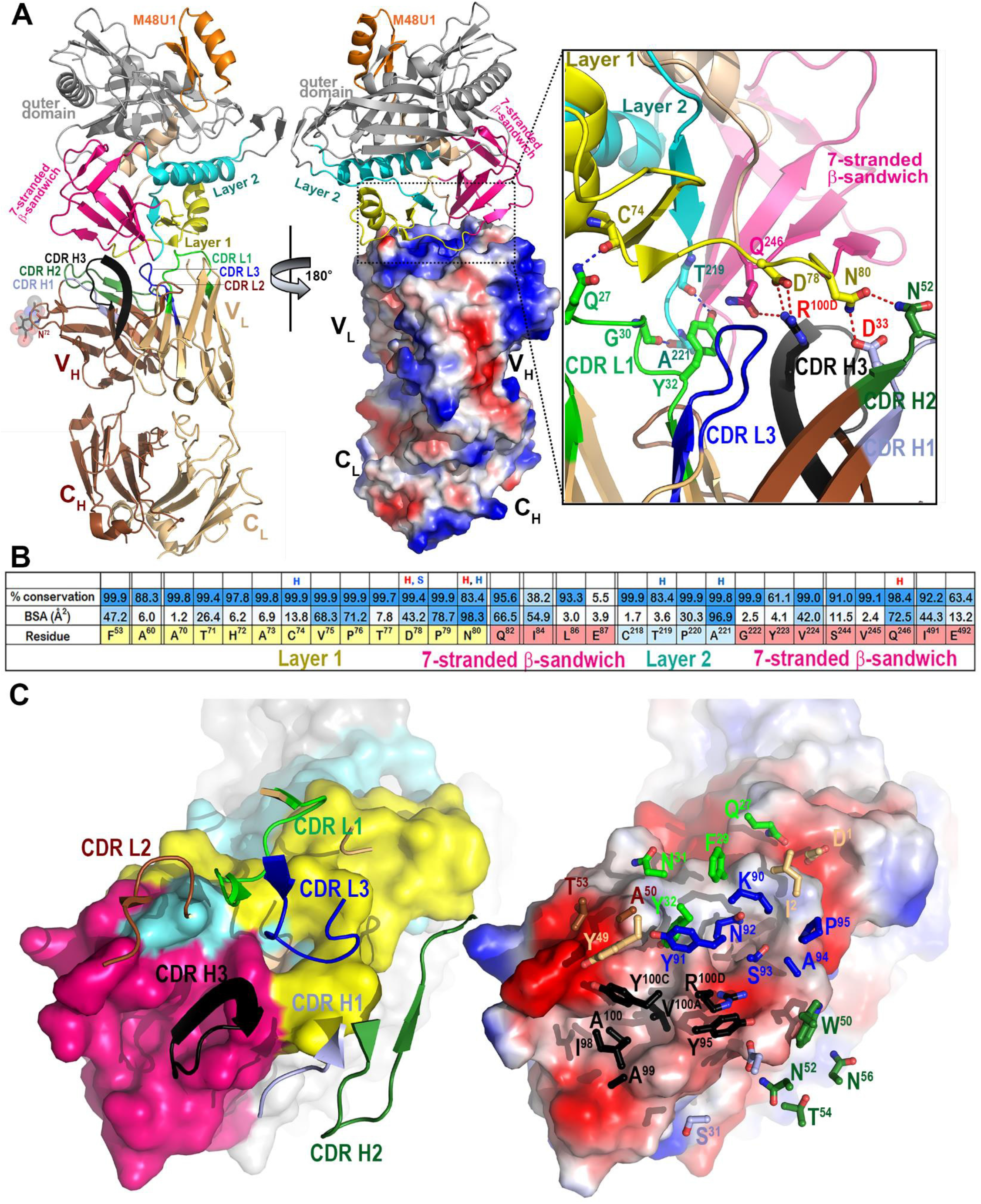
Crystal structure of the DH677.3 Fab-gp120_93TH057_ core_e_-M48U1 complex. **(A)** The overall structure of the complex is shown as a ribbon diagram (left) and with the molecular surface displayed over the Fab molecule (middle), colored based on electrostatic charge, red negative and blue positive. The gp120 outer domain is gray and inner domain colored to indicate inner domain mobile layer 1 (yellow), 2 (cyan), 3 (light orange) and the 7-stranded b-sandwich (magenta). Complementary determining regions (CDRs) are colored: CDR H1 (light blue), CDR H2 (dark green), CDR H3 (black), CRL1 (light green), CDR L2 (blue) and CDRL3 (brown). A blow-up view shows the network of hydrogen (H) bonds formed at the Fab-gp120 interface. H-bonds contributed by side chain and main chain atoms of gp120 residues are colored in magenta and blue, respectively. **(B)** Antibody buried surface area (BSA) and gp120 residues forming DH677.3 epitope are shaded in blue according to BSA (antibody) and percent conservation of gp120 residues (Env). gp120 main chain (blue) and side chain (red) hydrogen bonds (H) and salt bridges (S) are shown above the residue. **(C)** The DH677.3 Fab-gp120_93TH057_ core_e_ interface. CDRs are shown as ribbons (left) and balls-and-sticks of residues contributing the binding (right) over the gp120 core. The molecular surface of gp120 is colored as in **(A)** (left) and by electrostatic potential (right).

### Comparison of the DH677.3 mode of binding and epitope footprint to Cluster A prototype antibodies

Antigen complex structures of mAb A32 and N12-i3 (C11-like) [3, 6], antibodies isolated from HIV-1-infected individuals, confirm that DH677.3 recognized a unique epitope between the A32 and C11 antibody-binding sites involving Env epitope elements of both (**Fig 4**). While the A32 antibody epitope consists exclusively of gp120 mobile layers 1 and 2 (76% and 24% of gp120 BSA, respectively; **Table S4, Fig 4 B and C**), DH677.3 relies less on layers 1 and 2 (53% and 14% of gp120 BSA, respectively) and effectively utilizes the gp120 7-stranded β-sandwich (24% of gp120 BSA) (**Table S4, Fig 4 B and C**). The ability to recognize the 7-stranded β-sandwich renders DH677.3 similar to the C11-like antibody N12-i3, which almost exclusively depends on the β-sandwich for binding (94% of its total gp120 BSA; **Table S4, Fig 4 B and C**). Interestingly, N12-i3 and other C11-like antibodies require the N-terminus of gp120 for binding and recognize a unique gp120 conformation formed by docking of the gp120 N-terminus as an 8^th^ strand to the β-sandwich to form an 8-stranded-β-sandwich structure [6]. The DH677.3 complex crystals were obtained with gp120_93TH057_ core_e_ which lacks the N-terminus (Δ11 aa deletion) and therefore the direct judgment, based on structure, whether or not the 8^th^ strand is involved in binding was not possible (**Fig 3**). However, we were able to model the N/C-termini-gp120_93TH057_ core_e_ from the N12-i3 Fab complex structure (PDB code: 5W4L) to the DH677.3 Fab-gp120_93TH05_7 core_e_-M48U1 complex without any steric clashes (**Fig 4A, inset**). Both the conformation and orientation of CDR H1 and 2 of DH677.3 allowed easy access to the 8-stranded-β-sandwich structure and enabled contacts to the 8^th^ strand. These data indicated that DH677.3 is capable of accommodating both the 7 and 8-stranded-β-sandwich conformations of gp120 with effective contacts to the 8^th^ strand. Thus, the DH677.3 C1C2 antibody has a unique binding angle to the C1C2 region compared to C1C2 antibodies C11 and A32.

**Figure 4.**
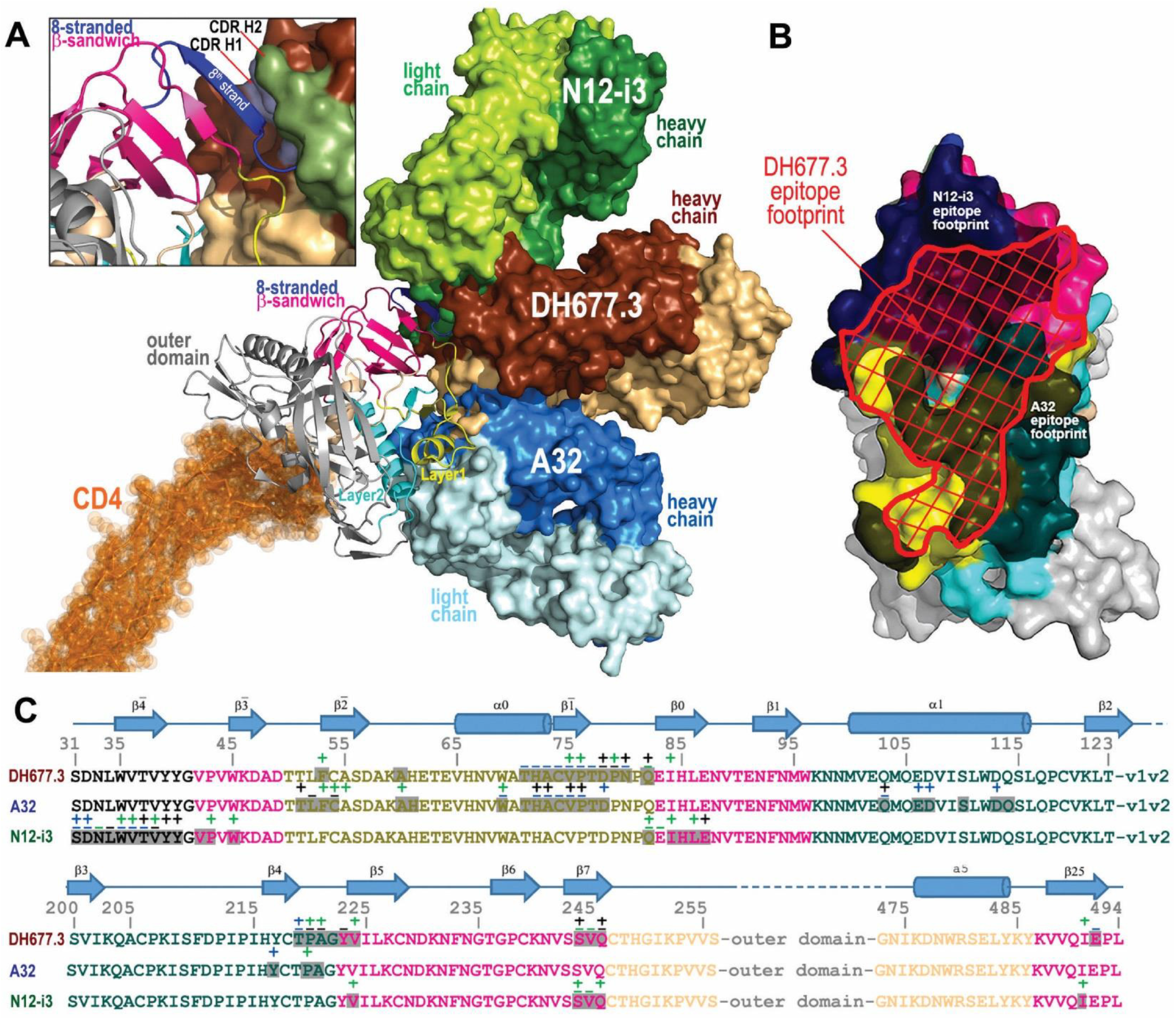
Recognition of HIV-1 Env by DH677.3 and other Cluster A antibodies. **(A)** The overlay of DH677.3 and Cluster A antibodies A32 and N12-i3 (C11-like) bound to the gp120 core. Crystals structures of the gp120 antigen in complex with the Fab of DH677.3, A32 (PDB code 4YC2) and N12-i3 (PDB code 5W4L) were superimposed based on gp120. The d1 and d2 domains of the target cell receptor CD4 was added to replace peptide mimetic M48U1 of the DH677.3 Fab-gp120_93TH057_ core_e_-M48U1 complex. Molecular surfaces are displayed over Fab molecules and colored in lighter and darker shades of brown, blue and green for the heavy and light chains of DH677.3, A32 and N12-i3, respectively. A blow up view shows details of the DH677.3 interaction with the 8-stranded β-sandwich of the gp120 inner domain. The 8^th^ strand (colored in blue) formed by the 11 N-terminal residues of gp120 in the N12-i3 bound conformation (PDB: 5W4L) was modeled into the DH677.3 Fab-gp120_93TH057_ core_e_-M48U1 complex. CHR H1 and 2 of DH677.3 are colored light blue and dark green, respectively. **(B)** and (C) Comparison of DH677.3, A32 and N12-i3 epitope footprints. In **(B)** the DH677.3 epitope footprint (shown in red) is plotted on the gp120 surface with layers colored as in Figure 1 with the A32 and N12-i2 epitope footprints shown in black. In **(C)** the DH677.3, A32 and N12-i3 gp120 contact residues are mapped onto the gp120 sequence. Side chain (+) and main chain (-) contact residues are colored green for hydrophobic, blue for hydrophilic and black for both as determined by a 5 Å cut off value over the corresponding sequence. Buried surface residues as determined by PISA are shaded. The DH677.3 epitope footprint overlays with the epitopes of both A32 and N12-i3.

### DH677 lineage antibodies mediate ADCC against CD4 downmodulated HIV-1 infected cells

During natural infection the HIV-1 accessory protein Nef downregulates CD4 expression on the surface of virus infected cells [12, 13]. Cell surface expressed CD4 facilitates the exposure of CD4i Env epitopes – like C1C2 - by binding to coexpressed cell surface Env [14]. The analyses of ADCC breadth was performed using target cells infected with IMCs containing the *Renilla* luciferase (LucR) reporter gene, which restricts Nef expression leading to incomplete CD4 downregulation [15]. Nevertheless, Vpu expression can compensate for Nef function and induce CD4 downregulation during the 72 hour incubation of the target cells before assays were performed. To exclude any possible impact of this technical aspect of IMCs with LucR on our ADCC results, full length IMCs (n=7) that do not contain a report gene were used to evaluate ADCC of the affinity matured RV305 C1C2-specific antibodies DH677.3 and DH677.4 and A32 [2], against target cells positive for intracellular p24 (p24+) and with downregulated CD4 (CD4-). As clade CRF01_AE possess a histidine at Env HXB2 position 375 that influences sensitivity to CD4i antibody binding and ADCC [16, 17] only clade B and clade C isolates were used.

When evaluating elimination of total p24+ cells no significant difference (Wilcoxon rank sum test; p > 0.05) in specific killing was noted among the three different antibodies (**Fig 5A**). However, when infected cells were separated into p24+CD4+ (**Fig 5B**) and p24+CD4-(**Fig 5C**) it was found that the RV305-boosted DH677.3 antibody was significantly better (Wilcoxon rank sum test p = 0.03) at mediating ADCC against p24+ CD4-infected cells (**Fig 5C**) when compared to A32. These data indicate that the DH677 clonal lineage epitope was more frequently exposed on Env conformers on the surface of IMC infected cells even in the context of CD4 downmodulation making this epitope a highly desirable NNAb vaccine target and important consideration in the setting of cure AIDS initiatives.

**Figure 5.**
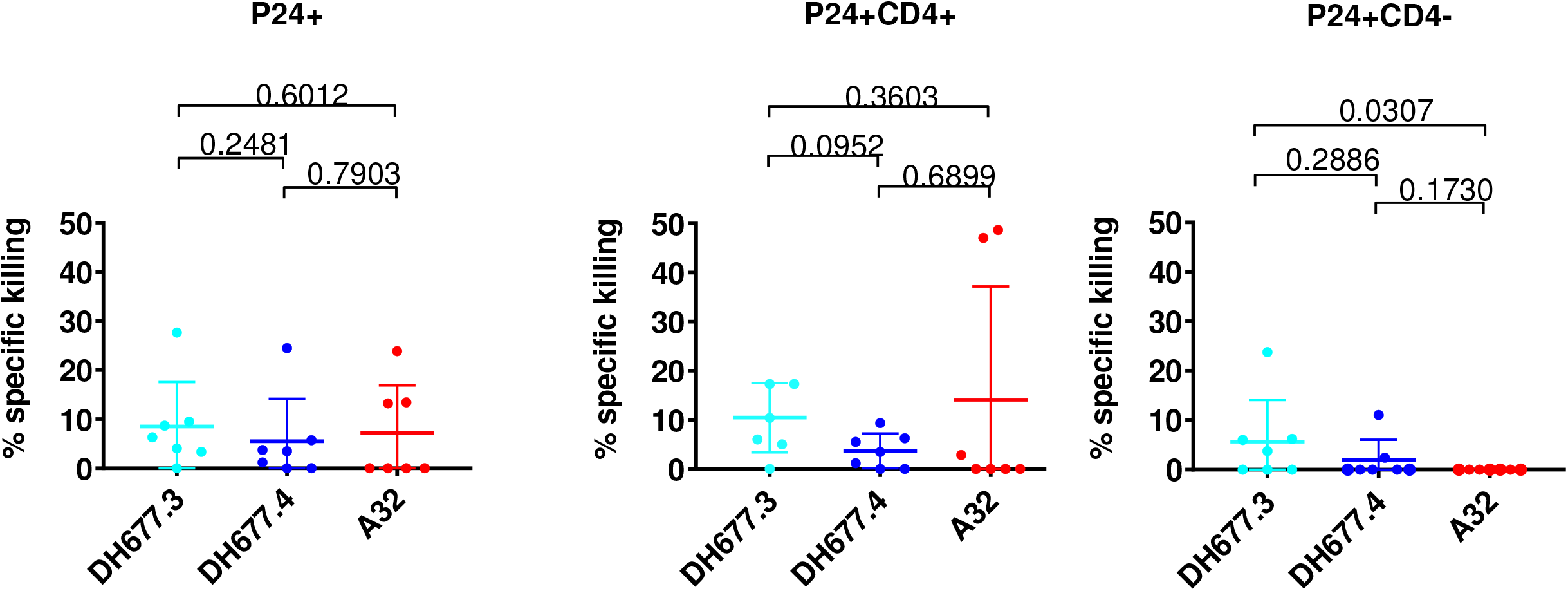
RV305 derived C1C2-specific antibody DH677.3 is significantly better than A32 at mediating antibody-dependent cellular cytotoxicity against CD4 down modulated infectious molecular clone infected cells. Cells were infected with clade B and clade C full length infectious molecular clones (IMC) that do not contain a reporter gene. Surface CD4 expression was analyzed by flow cytometry and p24 expression was measured in live/viable (A) all p24+ (B) p24+ CD4+ and (C) p24+ CD4-IMC infected cell populations. Data are shown with the mean and standard deviation.

## DISCUSSION

In this study it was found that late boosting of RV144 vaccinees increased C1C2-specific antibody V_H_ + V_L_ chain gene mutation frequency and increased clonal lineage specific ADCC breadth and potency (**Table 1, Fig 2**). Most RV305 derived antibodies had broader and more potent ADCC activity than the RV144 derived antibodies (**Table 1**) but V_H_ chain gene mutation frequency and the ADCC score did not directly correlate (**Fig S1**). While RV305 was necessary to mature C1C2-specific antibody responses, additional boosting with ALVAC/AIDSVAX B/E would not increase ADCC breadth and potency. Likely rationally designed sequential [18] boosting immunogens to select for critical mutations [19] that directionally affinity mature highly functional ALVAC/AIDSVAX B/E-induced C1C2-specific antibodies are needed.

Improving vaccine-induced NNAb effector function will also require more detailed immunological studies on the timing and frequency of boosting. In VAX003 (NCT00002441) and VAX004 (NCT00002441) trials frequent protein immunizations skewed Env-specific antibody subclass usage from the highly functional IgG3 to IgG4 [20–22]. The RV305 boosts that were studied here occurred several years (6-8yrs) later, unlike previous HIV-1 vaccine trials. Whether the boosting interval can be shortened without skewing antibody subclass usage is not known, but it is possible that boosting with long rest intervals (≥1-2 years) will be necessary.

The AIDSVAX B/E protein used for boosting in the RV144 and RV305 HIV-1 vaccine trial contained an N-terminal 11 amino acid deletion with important implications for NNAb induction. Previously it was shown that this modification enhanced exposure of the C1C2 region and V2 loop [9]. Here we show that this modification disrupts C11 – like antibody binding (**Fig 1**) but does creates a germline-targeting immunogen for DH677-like B cell lineages (**Fig S2**). Ligand crystal structure analysis found that DH677.3 recognized a unique C1C2 epitope that involves parts of epitope footprints of C1C2 Cluster A antibodies A32 and N12-i3 (C11-like) as well as new elements of the inner domain Layer 1 and the 7-stranded-β-sandwich (**Fig 3 and 4**). The DH677.3 epitope is positioned midway between the A32 and N12-i3 binding sites with most residues being highly conserved. Interestingly, DH677.3 binds at the edge of the gp120 inner domain 7-stranded β-sandwich and with layers 1 and 2 with a binding mode that allows it to accommodate the addition of the N-terminus as the 8^th^ strand to the 7-stranded-β-sandwich, a gp120 conformation emblematic of the late stages of HIV entry and recognized by C11 and C11-like antibodies [6]. Most likely this feature allows DH677.3 to recognize a broader range of Env targets, emerging in both the early (when A32 epitope becomes available) and late stage (when C11 epitope becomes available) of the viral entry process. Identification of a stage 2A of the HIV-1 Env expressed on the surface of infected cells in presence of the CD4 molecule or CD4 mimetics reiterate the importance of targeting these epitopes by vaccine induced responses as detected in our assays [23]. In addition, a model of DH677.3 in complex with gp120 antigen bound to a CD4 of a target/infected cell confirms that the recognition site and angle of approach position the DH677.3 IgG for easy access for effector cell recognition and Fc-effector complex formation (**Fig 4A**).

ADCC-mediating antibodies have been shown to reduce mother-to-child HIV-1 transmission [24–26],slow virus disease progression [26–28] and in RV144 correlated with reduced risk of infection in vaccine recipients with lower anti-Env plasma IgA responses [7]. Synergy between the RV144 C1C2 and V1V2 mAbs suggest a role for the C1C2 plasma responses that could not be directly identify by the correlates of protection study. Based on the data reported in this study and by Zoubchenok et al [16], it is clear that the magnitude of Env susceptibility to ADCC by the C1C2-specific Ab responses is not consistent as suggested by the conserved sequence of this region and varies according to the conformational stage of the HIV-1 envelope. That DH677.3 was better than A32 at mediating ADCC against HIV-1 clade B and C CD4 down-modulated cells (**Fig 5**) make this antibody an attractive candidate for targeting HIV-1 infected cells *in vivo* in the setting of HIV-1 infection. We have previously shown that the C1C2 antibody A32 when formulated as a bi-specific antibody can potently opsonize and kill HIV-1 infected CD4+ T cells [29]. Whether DH677.3-type of antibodies are superior to A32 for targeting virus-infected cells remains to be determined.

In summary, our data demonstrate that if the RV144 vaccine trial had been boosted, ADCC-mediating antibodies would have undergone affinity maturation for both ADCC potency and breadth of recognition of HIV-1-infected CD4+ T cells. Rationally designed subsequent boosting strategies to immunofocus IgG C1C2-specific response towards DH677-like antibody specificities may be one method by which to provide greater protection than observed in the RV144 HIV-1 vaccine trial.

## METHODS

### Ethics Statement

The RV305 clinical trial (NCT01435135) was a boost given to 162 RV144 clinical trial participants (NCT00223080) six-eight years after the conclusion of RV144 [30]. Donors used in this study were from groups boosted either with AIDSVAX B/E + ALVAC-HIV (vCP1521) (Group I) or AIDSVAX B/E alone (Group II). The RV305 clinical trial (NCT01435135) received approvals from Walter Reed Army Institute of Research, Thai Ministry of Public Health, Royal Thai Army Medical Department, Faculty of Tropical Medicine, Mahidol University, Chulalongkorn University Faculty of Medicine, and Siriraj Hospital. Written informed consent was obtained from all clinical trial participants. The Duke University Health System Institutional Review Board approved all human specimen handling.

### Antigen-specific single-cell sorting

1 × 10^7^ peripheral blood mononuclear cells (PBMCs) per vaccine-recipient were stained with AE.A244gp120Δ11 fluorescently labelled proteins and a human B cell flow cytometry panel. Viable antigen-specific B cells (AqVd-CD14-CD16-CD3-CD19+ IgD-) were single-cell sorted with a BD FACSAria II-SORP (BD Biosciences, Mountain View, CA) into 96 well PCR plates and stored at −80°C for RT-PCR.

### Single-cell reverse transcriptase PCR

Single B cell cDNA was generated with random hexamers using SSIII. The antibody variable heavy and light chain variable regions were PCR amplified using AmpliTaq360 Master Mix (Applied Biosystems). PCR products were purified (Qiagen, Valencia, CA) and sequenced by Genewiz. Gene rearrangements, clonal relatedness, unmutated common ancestors and intermediate ancestor inferences were made using Cloanalyst [31]. DH677 clonal lineage tree was generated using FigTree.

### Monoclonal antibody production

PCR-amplified heavy and light chain gene sequences were transiently expressed as previously described [32]. Ig containing cell culture supernatants were used for ELISA binding assays. For large scale expression, V_H_ and V_L_ chain genes were synthesized (V_H_ chain in the IgG1 4A backbone) and transformed into DH5α cells (GeneScript, Piscataway, NJ). Plasmids were expressed in Luria Broth, purified (Qiagen, Valencia, CA) and Expi293 cells were transfected using ExpiFectamine™ (Life Technologies, Carlsbad, CA) following the manufacturers protocol. After five days of incubation at 37°C 5% CO_2_ the Ig containing media was concentrated, purified with Protein A beads and the antibody buffer exchanged into PBS.

### Antibody binding and blocking assays

Direct ELISAs were performed as previously described [32]. Briefly, 384-well microplates were coated overnight with 30ng/well of protein. Antibodies were diluted and add for one hour. Binding was detected with an anti-IgG-HRP (Rockland) and developed with SureBlue Reserve TMB One Component (KPL). Plates were read on a plate reader (Molecular Devices) at 450nm. A32-blocking assays were performed by adding the RV305 antibodies followed by biotinylated A32 and detecting with streptavidin HRP.

### Neutralization assays

TZM-bl neutralization assays were performed in the Montefiori lab as previously described [33]. Data are reported as IC50 titers for antibodies.

### Infectious molecular clones (IMC)

The HIV-1 reporter virus used were replication-competent IMC designed to encode the *env* genes of CM235 (subtype A/E; GenBank No. AF259954.1), WITO (subtype B; GeneBank No. JN944948), 1086.c (subtype C; GeneBank No. FJ444395), TV-1 (subtype C; GeneBank No. HM215437), MW96.5 (subtype C; GeneBank No.), DU151 (subtype C; GeneBank No. DQ411851), DU422 (subtype C; GeneBank No. DQ411854) in *cis* within an Nef deficient isogenic backbone that expresses the *Renilla* luciferase reporter gene [34]. The subtype AE Env-IMC-LucR viruses used were the NL-LucR.T2A-AE.CM235-ecto (IMC_CM235_) (plasmid provided by Dr. Jerome Kim, US Military HIV Research Program), and clinical *env* isolates from the RV144 trial that were built on the 40061-LucR virus backbone. All the other IMCs were built using the original NL-LucR.T2A-ENV.ecto backbone as originally described by [35] Reporter virus stocks were generated by transfection of 293T cells with proviral IMC plasmid DNA, and virus titer was determined on TZM-bl cells for quality control [35]

### Infection of CEM.NKR_CCR5_ cell line with HIV-1 IMCs

CEM.NKR_CCR5_ cells were infected with HIV-1 IMCs as previously described [36]. Briefly, IMCs were titrated in order to achieve maximum expression within 48-72 hours post-infection as determined by detection of Luciferase activity and intra-cellular p24 expression. IMC infections were performed by incubation of the optimal dilution of virus with CEM.NKR_CCR5_ cells for 0.5 hour at 37°C and 5% CO_2_ in presence of DEAE-Dextran (7.5 μg/ml). The cells were subsequently resuspended at 0.5×10^6^/ml and cultured for 48-72 hours in complete medium containing 7.5μg/ml DEAE-Dextran. For each ADCC assay, we monitored the frequency infected target cells by intracellular p24 staining. Assays performed using infected target cells were considered reliable if cell viability was ≥60% and the percentage of viable p24+ target cells on assay day was ≥20%.

### Luciferase ADCC Assay

ADCC activity was determined by a luciferase (Luc)-based assay as previously described [8, 37] Briefly, CEM.NKR_CCR5_ cells (NIH AIDS Reagent Program, Division of AIDS, NIAID, NIH from Dr. Alexandra Trkola) [38] were used as targets after infection with the HIV-1 IMCs. PBMC obtained from a HIV-seronegative donor with the heterozygous 158F/V and 131H/R genotypes for Fc*γ*R3A and Fc*γ*R2A [39, 40], respectively, were used as a source of effector cells, and were used at an effector to target ratio of 30:1. Recombinant mAbs were tested across a range of concentrations using 5-fold serial dilutions starting at 50 μg/mL. The effector cells, target cells, and Ab dilutions were plated in opaque 96-well half area plates and were incubated for 6 hours at 37°C in 5% CO_2_. The final read-out was the luminescence intensity (relative light units, RLU) generated by the presence of residual intact target cells that have not been lysed by the effector population in the presence of ADCC-mediating mAb (ViviRen substrate, Promega, Madison, WI). The % of specific killing was calculated using the formula: percent specific killing = [(number of RLU of target and effector well – number of RLU of test well)/number of RLU of target and effector well] ×100. In this analysis, the RLU of the target plus effector wells represents spontaneous lysis in absence of any source of Ab. The ADCC endpoint concentration (EC), defined as the lowest concentration of mAb capable of mediating ADCC in our in vitro assay, was calculated by interpolation of the mAb concentration that intersected the positive cutoff of 15% specific killing. The RSV-specific mAb Palivizumab was used as a negative control.

### ADCC Score

Antibodies were tested across a range of concentrations using 5-fold serial dilutions starting at 50 μg/mL. Since the dilution curves are not monotonic due to prozone effect of mAbs, non-parametric area under the curve (AUC) was calculated using trapezoidal rule with activity less than 15% set to 0 %. For calculating a weighted average to obtain a score for ADCC activity explaining both potency and breadth of the mAbs, in this study we have used Principal Component Analysis (PCA) to compute an ADCC score. PCA is the most commonly used method to reduce the dimensionality of the data set [41]. It uses Eigen vector decomposition of the correlation matrix of the variables, where each variable is represented by a viral isolate. Most of the shared variance of the correlations of ADCC AUC is explained by first principal component (PC1) [42]. Ideally, one would want to explain 70% of the variance but should not be at the expense of adding principal components with an Eigenvalue less than 1 [43].

In this study we a panel of 7 HIV isolates was tested which implies that our data set has seven dimensions. ADCC activity was measured as AUC. In our analysis PC1 and PC2 have Eigen values above 1 and together account for 80.57% variance (**Table S5**). Scores obtained from the first Principal Component can be interpreted as weighted average of the 7 isolates that would account for both potency as well as breadth of the mAbs [43]. Higher PC1 score would mean that mAb has a higher breadth as well as potency for ADCC activity. To calculate the ADCC score, the standardized AUC value for each monoclonal antibody is calculated for each viral isolate, multiplied by factor loading of the given viral isolate and then these products are added together. Standardized AUC values imply zero mean and unit standard deviation. The AUC values below the value of mean AUC will result in negative PC1 scores.

### Infection of primary cells with HIV-1 IMCs

Infectious molecular clones encoding the full-length transmitted/founder sequence of seven individuals infected with either subtype B or C viruses from the CHAVI acute infection cohort (CH77, CH264, CH0470, CH042, CH185, CH162 and CH236) were constructed as previously described [44, 45] and used to infect primary CD4+ cells. To infect cells, cryopreserved peripheral blood mononuclear cells (PBMCs) were thawed and stimulated in R20 media (RPMI media (Invitrogen) with 20% Fetal Bovine Serum (Gemini Bioproducts), 2mM L-glutamine (Invitrogen), 50 U /mL penicillin (Invitrogen), and 50 μg/mL Gentamicin (Invitrogen)) supplemented with IL-2 (30U/mL, Proleukin), anti-CD3 (25ng/mL clone OKT-3, Invitrogen) and anti-CD28 (25ng/mL, BD Biosciences) antibodies for 72 hours at 37°C in 5% CO_2_. CD8 cells were depleted from the PBMCs using CD8 microbeads (Miltenyi Biotec, Germany) according to the Manufacturer’s instructions and 1.5 × 10^6^ cells were infected using 1 mL virus supernatant by spinoculation (1125 × g) for 2 hours at 20 °C. After spinoculation, 2 mL of R20 supplemented with IL-2 was added to each infection and infections were left for 72 hours. Infected cells were used if viability was >70% and more than 5% of cells were p24+.

### Infected Cell Elimination Assay

HIV-1-infected or mock-infected CD8-depleted PBMCs cells were used as targets and autologous cryo-preserved PBMCs rested overnight in R10 supplemented with 10ng/ml of IL-15 (Miltenyi Biotec) were used as a source of effector cells. Infected and uninfected target cells were labelled with a fluorescent target-cell marker (TFL4; OncoImmunin) and a viability marker (NFL1; OncoImmunin) for 15 min at 37 °C, as specified by manufacturer. The labeling of the target cells with these two markers allowed to clearly identify only the live viable cells in our gating strategy and exclude artifacts related to the presence of dead cells staining. Cells were washed in R10 and adjusted to a concentration of 0.2×10^6^ cells/mL. PBMCs were then added to target cells at an effector/target ratio of 30:1 (6 × 10^6^ cells/mL). The target/effector cell suspension was plated in V-bottom 96-well plates and co-cultured with 10 μg/mL of each mAb. Co-cultures were incubated for 6 h at 37 °C in 5% CO_2_. After the incubation period, cells were washed and stained with anti-CD4-PerCP-Cy5.5 (eBioscience, clone OKT4) at a final dilution of 1:40 in the dark for 20 min at room temperature (RT). Cells were then washed, resuspended in 100 μL/well Cytofix/Cytoperm (BD Biosciences), incubated in the dark for 20 min at 4 °C, washed in 1x Cytoperm wash solution (BD Biosciences) and co-incubated with anti-p24 antibody (clone KC57-RD1; Beckman Coulter) to a final dilution of 1:100, and incubated in the dark for 25 min at 4 °C. Cells were washed three times with Cytoperm wash solution and resuspended in 125 μL PBS-1% paraformaldehyde. The samples were acquired within 24 h using a BD Fortessa cytometer. The appropriate compensation beads were used to compensate the spill over signal for the four fluorophores. Data analysis was performed using FlowJo 9.6.6 software (TreeStar). Mock-infected cells were used to appropriately position live cell p24+/− and CD4+/− gates.

Specific killing was determined by the reduction in % of viable p24+ cells in the presence of mAbs after taking into consideration non-specific killing, and was calculated as:

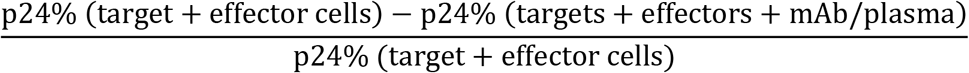

CH65 (an anti-influenza monoclonal antibody, kindly provided by Dr. Moody) was used as negative control. To remove background signal, the highest value of percent specific killing induced by CH65 was subtracted from the calculated reduction in % of p24+ cells and then negative values were rounded to 0%.

### Surface plasmon resonance

The binding and kinetic rates measurement of gp120 proteins against RV305 mAbs were obtained by surface plasmon resonance (SPR) using the Biacore 3000 instrument (GE Healthcare). SPR measurements were performed using a CM5 sensor chip with anti-human IgG Fc antibody directly immobilized to a level of 9000-11000RU (response unit). Antibodies were then captured at 5ul/min for 60s to a level of 100-300RU. For binding analyses, the gp120 proteins were diluted to approximately 1000nM in PBS and injected over the captured antibodies for 3 minutes at 30ul/min. For kinetics measurements, the gp120 proteins were diluted from 5-750nM and injected using a high performance kinetics injection for 5 minutes at 50uL/min. This was followed by a dissociation period of 600s and surface regeneration with Glycine pH2.0 for 20s. Results were analyzed using the Biacore BiaEvaluation Software (GE Healthcare). Negative control antibody (Ab82) and blank buffer binding were used for double reference subtraction to account for non-specific protein binding and signal drift. Subsequent curve fitting analysis was performed using a 1:1 Langmuir model with a local Rmax and the reported rate constants are representative of two measurements.

### Protein preparation and complex crystallization

DH677.3 Fab alone was grown and crystalized at concentration ~10 mg/ml. The structure was solved by molecular replacement with PDB ID 3QEG in space group P21 to a resolution of 2.6 Å. Clade A/E 93TH057 gp120 core_e_, (gp120_93TH057_ core_e_, residues 42-492 (Hxbc2 numbering)), lacking the V1, V2 and V3 variable loops and containing a H375S mutation to allow binding of the CD4 mimetic M48U1 [46] was used to obtain crystals of DH677.3 Fab-antigen complex. gp120_93TH057_ core_e_ was prepared and purified as described in [3]. Deglycosylated gp120_93TH057_ core_e_ was first mixed with CD4 mimetic peptide M48U1 at a molar ratio of 1:1.5 and purified through gel filtration chromatography using a Superdex 200 16/60 column (GE Healthcare, Piscataway, NJ). After concentration, the gp120_93TH057_ core_e_-M48U1 complex was mixed with a 20% molar excess of DH677.3 Fab and passed again through the gel filtration column equilibrated with 5 mM Tris-HCl buffer pH 7.2 and 100 mM ammonium acetate. The purified complex was concentrated to ~10 mg/ml for crystallization experiments. The structure was solved by molecular replacement using the DH677.3 Fab and PDB ID 3TGT as searching models in space group P1 to a resolution 3.0 Å. The final R_factor_/R_free_ (%) for the Fab structure is 19.9/26.1 and the final R_factor_/R_free_ for the complex is 21.4/27.4 (**Table S3**). The PDB IDs for the deposited structures are 6MFJ and 6MFP respectively. In each case the asymmetric unit of the crystal contained two almost identical copies of Fab or the Fab-gp120_93TH057_ core_e_ complex (**Fig S2**)

### Crystallization and Data collection

Initial crystal screens were done in vapor-diffusion hanging drop trials using commercially available sparse matrix crystallization screens from Hampton Research (Index), Emerald BioSystems (Precipitant Wizard Screen) and Molecular Dimensions (Proplex and Macrosol Screens). The screens were monitored periodically for protein crystals. Conditions that produced crystals were then further optimized to produce crystals suitable for data collection. DH677.3 Fab crystals were grown from 20% PEG 3000, 100 mM HEPES pH 7.5, and 200 mM sodium chloride. DH677.3 complex crystals were grown from 25% PEG 4000 and 100 mM MES pH 5.5. Crystals were briefly soaked in crystallization solution plus 20% MPD before being flash frozen in liquid nitrogen prior to data collection.

### Data collection and structure solution

Diffraction data were collected at the Stanford Synchrotron Radiation Light Source (SSRL) at beam line BL12-2 equipped with a Dectris Pilatus area detector. All data were processed and reduced with HKL2000 [47]. Structures were solved by molecular replacement with Phaser [48] from the CCP4 suite [49]. The DH677.3 Fab structure was solved based on the coordinates of the N12-i2 Fab (PDB: 3QEG), and the DH677.3 complex was then solved with coordinates from the DH677.3 Fab model, gp120 (PDB: 3TGT), and M48U1 (PDB: 4JZW). Refinement was carried out with Refmac [50] and/or Phenix [51]. Refinement was coupled with manual refitting and rebuilding with COOT [52]. Data collection and refinement statistics are shown in **Table 1**.

### Structure validation and analysis

The quality of the final refined models was monitored using the program MolProbity [53]. Structural alignments were performed using the program lsqkab from the CCP4 suite [49]. The PISA [54] webserver was used to determine contact surfaces and residues. All illustrations were prepared with the PyMol Molecular Graphic suite (http://pymol.org) (DeLano Scientific, San Carlos, CA, USA). Conservation of the DH677.3 epitope was calculated using the HIV Sequence Database Compendium (https://www.hiv.lanl.gov/content/sequence/HIV/COMPENDIUM/compendium.html) comparing gp120 residues relative to Clade B Hxbc2. Only unique sequences in the database having an equivalent residue at each position were included in the calculated percentage representing approximately 32,000 sequences on average.

### Statistical Methods

For luciferase based ADCC assay background correction was performed by subtracting the highest value of percent specific killing induced by CH65 and then rounding off the negative values to zero. _In order to assess if two groups have different response pairwise comparisons between groups was conducted using Wilcoxon rank sum test. Statistical analysis was performed using SAS software (SAS Institute Inc., Cary, N.C.).

---

**Figure S1. RV305 C1C2-specific antibody ADCC score inversely correlate with antibody mutation frequency.** Correlation between RV305 antibody heavy chain gene mutation frequency (% nucleotide; Cloanalyst) [31] and ADCC score (see methods) was calculated with SAS (Spearman Correlation = −0.5599; p value = 0.0127).

**Figure S2. RV305 boosting increased affinity of the C1C2-specific RV144 derived DH677 memory B cell clonal lineage to the AIDSVAX B/E proteins**. DH677.1 antibody was isolated by antigen-specific single-cell sorting PBMC from the RV144 vaccine trial. DH677.2, DH677.3 and DH677.4 were isolated by antigen-specific singlecell sorting PBMC collected after the second boost AIDSVAX B/E (RV305 Group II) given in RV305 (~7yrs later). The intermediate ancestors and unmutated common ancestor was inferred using Clonalayst [31]. Recombinantly expressed antibodies were assayed by surface plasmon resonance for binding to the AIDSVAX B/E proteins – AE.A244g120Δ11 + B.MNg120Δ11- and to full length AE.A244gp120.

**Figure S3. Boosting in RV305 increased DH677 B cell clonal lineage cross-clade antibody-dependent cellular cytotoxicity breadth and potency**. Recombinantly expressed DH677 clonal lineage members were assayed for antibody-dependent cellular cytotoxicity (ADCC) against AE.CM235, B.WITO, C.TV-1, C.MW965, C.1086C, C. DU151 and C.DU422 Renilla luciferase reporter gene infectious molecular clone infected cells. Data are shown as radar plots with an ADCC score (see methods) that accounts for ADCC breadth and potency.

**Figure S4. Comparison of the two copies of the DH677.3 Fab-gp120_93TH057_ core_e_-M48U1 complex and the two Fab copies in the apo Fab structure from the asymmetric unit of crystals.** (A) The root mean square deviation (RMSD) between complex copies is 0.946 Å for main chain residues. **(B)** The RMSD between the Fab copies in the apo Fab structure is 0.540 Å for main chain residues. **(C)** Comparison of the free and bound DH677.3 Fab. The α-carbon backbone diagram of superposition of the structures of DH677.3 Fab alone (dark cyan-heavy chain and light cyan-light chain) and N5-i5 Fab bound to CD4-triggered gp120 (dark brown-heavy chain and light brown-light chain). The average RMSD between free and bound Fabs is 0.818 Å for main chain residues.

**Figure S5. Antibody contact residues**. mAb side chain (+) and main chain (-) contact residues colored green for hydrophobic, blue for hydrophilic and black for both as determined by a 5 Å cut off value over the corresponding sequence. CDRs are colored as in Figure 1 and buried surface residues as determined by PISA are shaded.

**Table S1. Immunogenetics of non-neutralizing A32-blocking RV305 C1C2-specific antibodies**. RT-PCR amplified variable heavy and variable light chain genes were Sanger sequenced (Genewiz) and analyzed with Cloanalyst [31].

**Table S2. A32-blocking antibodies do not neutralize HIV-1**. (A) Recombinantly expressed antibodies were assayed in the TZM-bl neutralization assay against autologous and heterologous Tier 1 and Tier 2 isolates. No neutralization was detected. Data are shown as EC50 (μg/mL)

**Table S3. DH677.3 structural data collection and refinement statistics**

**Table S4. Details of the DH677.3, A32, and N12-i3 interfaces based on the DH677.3-gp120_93TH057_ core_e_-M48U1, A32 Fab-ID2_93TH057_, and N12-i3 Fab-gp120_93TH057_core_e_+N/C-M48U1 structures as** calculated by the EBI PISA server (http://www.ebi.ac.uk/msd-srv/prot_int/cgi-bin/piserver). The two copies in the asymmetric unit of the DH677.3, A32, and N12-i3 complexes are averaged in the table.

## ACKNOWLEDGEMENTS

The authors would like to acknowledge the Duke Human Vaccine Institute Flow Cytometry Facility (Durham, NC), Duke Human Vaccine Institute Viral Genetic Analysis Facility (Durham, NC) and the following individuals for their expert technical assistance: flow cytometry – Derek Cain, Patrice McDermott and Dawn Jones Marshall, conjugated antigens – Lawrence Armand, transient transfections – Andrew Foulger, Erika Dunford and Kedamawit Tilahun, ELISA – Rob Parks, Callie Vivian and Maggie Barr, Antibody Expression – Giovanna Hernandez, Esther Lee, Emily Machiele and Rachel Reed, Neutralization Assays – Amanda Eaton, Celia C. LeBranche, Peter Gao, Kelli Greene and Hongmei Gao, Biolayer Interferometry – Kara Anasti. From project management – Cynthia Nagle and Kelly Soderberg. We also thank all of the RV144 and RV305 clinical trial team members and participants. This work was primarily supported by a Collaboration for AIDS Vaccine Discovery Grant OPP1114721 from the Bill & Melinda Gates Foundation to BFH, and by NIH grants NIAID R01 AI116274 and R01 AI129769 to MP, NIAID P01 AI120756 to GT, and a Henry M. Jackson Foundation for the Advancement of Military Medicine #829295 grant to BFH.

## AUTHOR CONTRIBUTION

Conceptualization, D.E., B.F.H., J.P., M.P., G.F.; Methodology, D.E., B.F.H., J.P., G.F.; Software, K.W., T.B.K.; Validation, D.E., J.P., S.J., M.P., G.F.; Formal Analysis, S.J., K.W.; Investigation, D.E., J.P., K.L., W.D.T., B.Y., D.M.; Resources, R.J.O., S.V., J.K., K.K., P.P., S.N., F.S., J.T., S.P.; Data Curation, D.E., J.P., K.W., D.M., S.J., S.M.A., D. C.M., M.P., G.F.; Writing – Original Draft, D.E., S.J., M.P., G.F.; Writing – Review & Editing, D.E., J.P., W.D.T., B.Y., S.J., S.V., S.P., G.D.T., M.A.M., M.P., B.F.H., G.F.; Visualization, D.E., J.P., S.J., G.F.; Supervision, D.E., J.P., W.D.T., K.O.S., D.C.M., M.P., B.F.H., G.F.; Project Administration; D.E., M.P., G.F., B.F.H.; Funding Acquisition, M.P., G.D.T., B.F.H., G.F.

## DISCLAIMER

The views expressed are those of the authors and should not be construed to represent the positions of the Uniformed Services University, U.S. Army, Department of Defense or the Department of Health and Human Services. The investigators have adhered to the policies for protection of human subjects as prescribed in AR-70.

